# Zebrafish Nanog is not required in embryonic cells

**DOI:** 10.1101/091504

**Authors:** James A. Gagnon, Kamal Obbad, Alexander F. Schier

**Affiliations:** Department of Molecular and Cellular Biology, Harvard University, Cambridge MA, USA; Center for Brain Science, Harvard University, Cambridge MA, USA; The Broad Institute of Harvard and MIT, Cambridge MA, USA; FAS Center for Systems Biology, Harvard University, Cambridge MA, USA

**Keywords:** Nanog, zebrafish, maternal-to-zygotic transition, MZT, yolk syncytial layer, YSL, zygotic genome activation, ZGA, embryogenesis

## Abstract

The study of *nanog* mutants reveals that Nanog is required only for extraembryonic tissue development, not in embryonic cells.

**ABSTRACT:** The role of the zebrafish transcription factor Nanog has been controversial. It has been suggested that Nanog is primarily required for the formation of the extraembryonic yolk syncytial layer (YSL) and only indirectly regulates gene expression in embryonic cells. By contrast, a more recent study has proposed that Nanog directly regulates transcription in embryonic cells during zygotic genome activation. To clarify the roles of Nanog, we performed a detailed analysis of zebrafish *nanog* mutants. While zygotic *nanog* mutants survive to adulthood, maternal-zygotic and maternal mutants exhibit developmental arrest at the blastula stage. In the absence of Nanog, the YSL fails to form and embryonic tissue detaches from the yolk. Zygotic transcription of a subset of embryonic genes is affected in *nanog* mutants but both the YSL and embryonic phenotype can be rescued by providing *nanog* mRNA in YSL precursors. Notably, *nanog* mutant cells transplanted into wild-type hosts proliferate and contribute to embryonic tissues from all germ layers. These results indicate that zebrafish Nanog is necessary for YSL formation but is not directly required for embryonic cell differentiation.

## INTRODUCTION

The transcription factor Nanog is part of the core circuitry that regulates mammalian pluripotency (reviewed in Theunissen & Jaenisch 2014). In vitro studies have shown that removal of Nanog triggers differentiation of mouse and human embryonic stem cells (Mitsui et al. 2003; Chambers et al. 2007; Hyslop et al. 2005; Loh et al. 2006). However, a subset of *nanog* mutant mouse embryonic stem cells are able to self-renew (Chambers et al. 2007). In vivo studies have revealed that *nanog* is required for inner cell mass pluripotency and epiblast development (Mitsui et al. 2003). However, in chimeras with wild-type cells, *nanog* mutant cells can give rise to tissues from all germ layers (Chambers et al. 2007). Thus, mouse Nanog is involved in but not absolutely required for the maintenance of the pluripotent state.

The roles of zebrafish Nanog in pluripotency and differentiation are less well understood. Xu et al. (2012) reported that *nanog* was provided maternally and present in all embryonic and extraembryonic cells. Morpholino-mediated knockdown of *nanog* resulted in developmental arrest prior to gastrulation. Nanog morphants lacked the YSL, the extraembryonic tissue that attaches the embryo to the yolk and generates Nodal and BMP signals that pattern mesendoderm (Xu et al. 2012; Carvalho & Heisenberg 2010; Hong et al. 2011; Chen & Kimelman 2000; Mizuno et al. 1996). Gene expression analysis in *nanog* morphants revealed the absence of YSL markers such as *mxtx2* and the misregulation of hundreds of embryonic genes, including Nodal and its target genes. Injecting *mxtx2* mRNA into YSL precursors of *nanog* morphants partially rescued YSL formation and the expression of Nodal and several of its target genes. Although no cell autonomy data was shown to determine whether Nanog was required in embryonic cells, the study suggested that Nanog primarily regulates the formation of the YSL (Xu et al. 2012).

Two subsequent studies analyzed potential roles of zebrafish Nanog in embryonic cells (Lee et al. 2013; Perez-Camps et al. 2016). Lee et al. (2013) defined a set of genes expressed at the maternal-to-zygotic transition (MZT) whose expression was reduced in *nanog* morphants. Chromatin immunoprecipitation experiments suggested that many of these genes were direct targets of Nanog (Lee et al. 2013; Xu et al. 2012; Leichsenring et al. 2013; Bogdanovic et al. 2012). Based on the reduced expression of genes in morphants and the Nanog binding data, the study concluded that Nanog, along with Pou5f1 and SoxB1, was involved in the first wave of zygotic transcription in embryonic cells. Subsequent reviews have interpreted these results to conclude that Nanog is directly required for zygotic genome activation in embryonic cells (Langley et al. 2014; Paranjpe & Veenstra 2015; Onichtchouk & Driever 2016; Lee et al. 2014), even though the majority of zygotic genes are activated in *nanog* morphants (Lee et al. 2013; Xu et al. 2012). Perez-Camps et al. (2016) reported that morpholino knockdown of *nanog* caused defects in BMP signaling and target gene expression, and suggested that Nanog acts to promote ventral cell fate specification. Surprisingly, neither study (Perez-Camps et al. 2016; Lee et al. 2013) mentioned the YSL phenotype of *nanog* morphants (Xu et al. 2012) or tested for direct roles of Nanog in embryonic cells.

Here we clarify the embryonic and extraembryonic requirements for Nanog using tissue-specific rescue and chimera analysis. Our results indicate that zebrafish Nanog is only essential for YSL formation and not directly required for embryonic cell differentiation.

## RESULTS

### Generation of *nanog* mutants

The interpretation of morpholino experiments can be complicated by potential partial loss-of-function phenotypes and the short half-life of morpholinos. To avoid these confounds in our studies of Nanog, we generated *nanog* mutants using TALENs (Carroll 2014). We isolated an allele containing a 7 bp deletion predicted to cause a frameshift and premature termination codon before the homeodomain required for DNA binding (**Figure 1A**). The mutant *nanog* mRNA was not detectable at sphere stage (4 hours post-fertilization [hpf]), presumably due to nonsense-mediated decay (**Figure 1B**). Homozygous zygotic *nanog* (Z*nanog*) mutant embryos showed no phenotypic defects and could be raised to fertile adults. By contrast, maternal-zygotic *nanog* mutants (MZ*nanog*) and maternal-only *nanog* mutants (M*nanog*) arrested at sphere stage, did not undergo normal epiboly (**Figure 1C**), and died when embryonic tissue detached from the yolk. The defects observed in MZ*nanog* embryos were rescued by injection of *nanog* mRNA at the 1-cell stage (**Figure 1D**). Rescued embryos could be raised to fertile adults, establishing that the observed phenotype is solely due to disruption of the *nanog* gene. The *nanog* mutant phenotype strongly resembles the previously reported *nanog* morphant phenotype (Xu et al. 2012). Since M*nanog* and MZ*nanog* had identical mutant phenotypes, while Z*nanog* was viable, we conclude that maternal, but not zygotic, Nanog has essential roles during embryogenesis.

**Figure 1.**
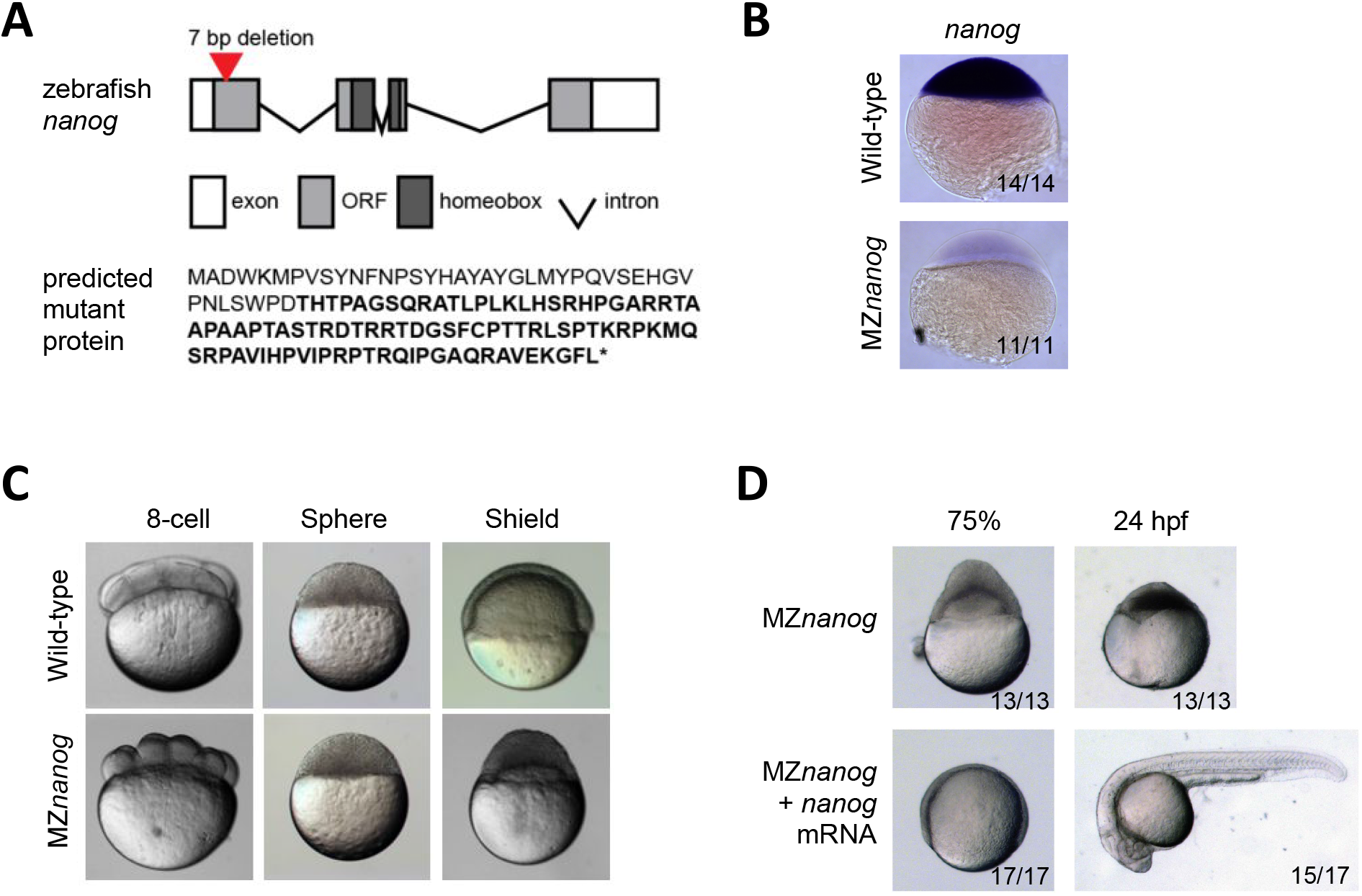
Generation and phenotype of MZ*nanog* mutants. **A.** (top) TALENs were used to generate a seven basepair (bp) deletion within the first exon of *nanog*. Exons, introns, the open reading frame (ORF) and the homeobox domain are indicated. (bottom) The predicted mutant protein sequence, with intact amino acids (40 of 384 total) in normal font and frameshifted amino acids in bold font. Asterisk indicates a premature stop codon. **B.** In situ hybridization for *nanog* expression in wild-type and MZ*nanog* embryos at sphere stage. **C.** Wild-type and MZ*nanog* embryos imaged at 8-cell (1.25 hpf [hours post-fertilization]), sphere (4 hpf), and shield (6.5 hpf) stages. Epiboly defects in MZ*nanog* are apparent at shield stage. **D.** MZ*nanog* embryos are shown at 75% epiboly (8 hpf) stage or at 24 hpf, either uninjected or injected with 5 pg *nanog* mRNA at the 1-cell stage.

### *Nanog* is required for formation of the YSL

*Nanog* morphants are impaired in YSL development and the expression of YSL markers (Xu et al. 2012). Three lines of evidence indicate that the YSL is also absent in MZ*nanog* mutants. First, MZ*nanog* embryos lacked expression of the YSL determinant *mxtx2* at sphere stage (**Figure 2A**). Second, expression of other YSL marker genes such as *slc26a1, gata3,* and *hnf4a* (Xu et al. 2012) was decreased or absent as determined by RT-qPCR (**Figure 2B**). Third, RNAseq experiments showed that the expression of many genes expressed in the YSL (Xu et al. 2012) was reduced in MZ*nanog* mutants at shield stage (6.5 hpf; **Figure 2C**). Expression of many of these genes was not completely eliminated, likely due to their expression in embryonic cells. Compared to zygotically expressed housekeeping genes **(Figure 2D)**, YSL genes were significantly downregulated in MZ*nanog* embryos (p=7.5x10^-6^, **Figure 2E)**. These results confirm and extend previous morphant studies that concluded that Nanog is required for YSL specification (Xu et al. 2012).

**Figure 2.**
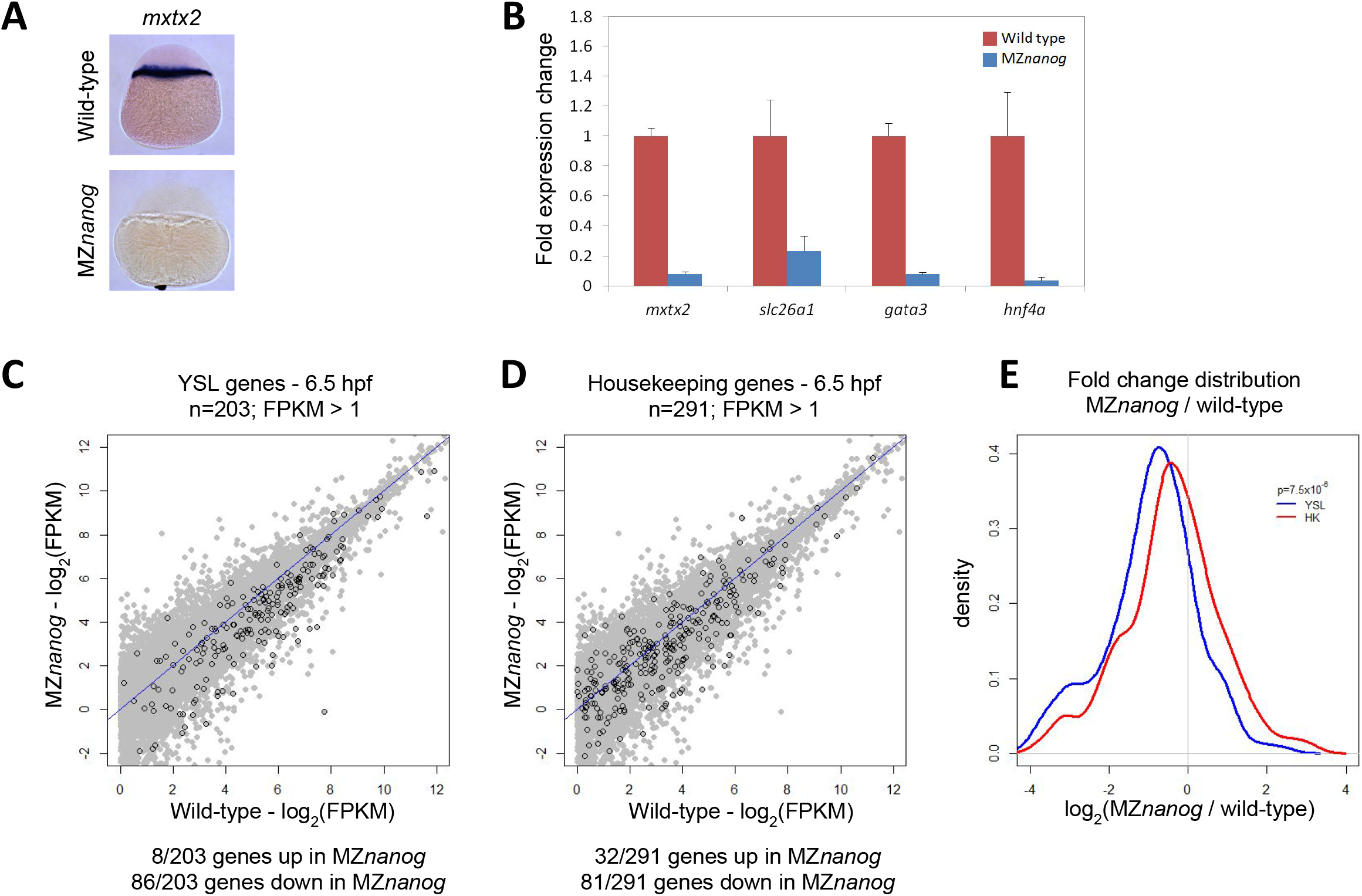
Absence of the YSL in MZ*nanog* embryos. **A.** In situ hybridization for *mxtx2* expression in wild-type and MZ*nanog* embryos at sphere stage. **B.** Fold expression change for *mxtx2, slc26a1, gata3* and *hnf4a* comparing wild-type and MZ*nanog* embryos at sphere stage using RT-qPCR. Error bars show standard deviation for three technical replicates. **C.** Differential expression of genes expressed in the YSL, comparing wild-type and MZ*nanog* at shield stage (6.5 hpf) using RNAseq. All genes are in grey, YSL expressed genes are in black and were previously defined (Xu et al. 2012), and filtered for zygotic expression (Rabani et al. 2014). **D.** Differential expression of zygotically expressed housekeeping genes (Lee et al. 2013), comparing wild-type and MZ*nanog* at shield stage (6.5 hpf) using RNAseq. All genes are in grey, housekeeping genes are in black. Only those genes with wild type mRNA expression > 1 FPKM are plotted. Genes are categorized as up- or down-regulated if their expression in MZ*nanog* differs more than two-fold from that in wild-type. **E.** Distribution of fold changes derived from RNAseq data comparing wild-type and MZ*nanog* embryos for all YSL and housekeeping (HK) genes using a kernel density estimation. The displayed p-value comparing these two sets was calculated using a Student’s two-sided t-test.

### *Nanog* regulates the expression of a subset of early zygotic genes

*Nanog* morphants were reported to display widespread reduction in zygotic gene expression (Lee et al. 2013). To determine if similar defects are found in MZ*nanog* mutants, we used in situ hybridization, RT-qPCR and RNAseq. By in situ hybridization and RT-qPCR, the expression of several early zygotic genes was decreased in MZ*nanog* (**Figure 3A and B**). RNAseq experiments revealed that the expression of many of the previously defined early zygotic genes (Lee et al. 2013) was reduced in MZ*nanog* embryos at sphere stage (**Figure 3C**). When compared to maternally provided mRNAs, which are largely unaffected **(Figure 3D)**, the levels of early zygotic RNAs are significantly reduced in MZ*nanog* mutants (p=0.00307, **Figure 3E).** The expression of 79/251 early zygotic genes was reduced >2-fold in MZ*nanog* mutants. Many of these genes are part of developmental signaling pathways (**Table S1**). 68 of the 79 genes reduced in MZ*nanog* mutants were also significantly decreased in the *nanog* morphants (Lee et al. 2013). These results indicate that *nanog* mutants and morphants have similar gene expression defects.

**Figure 3.**
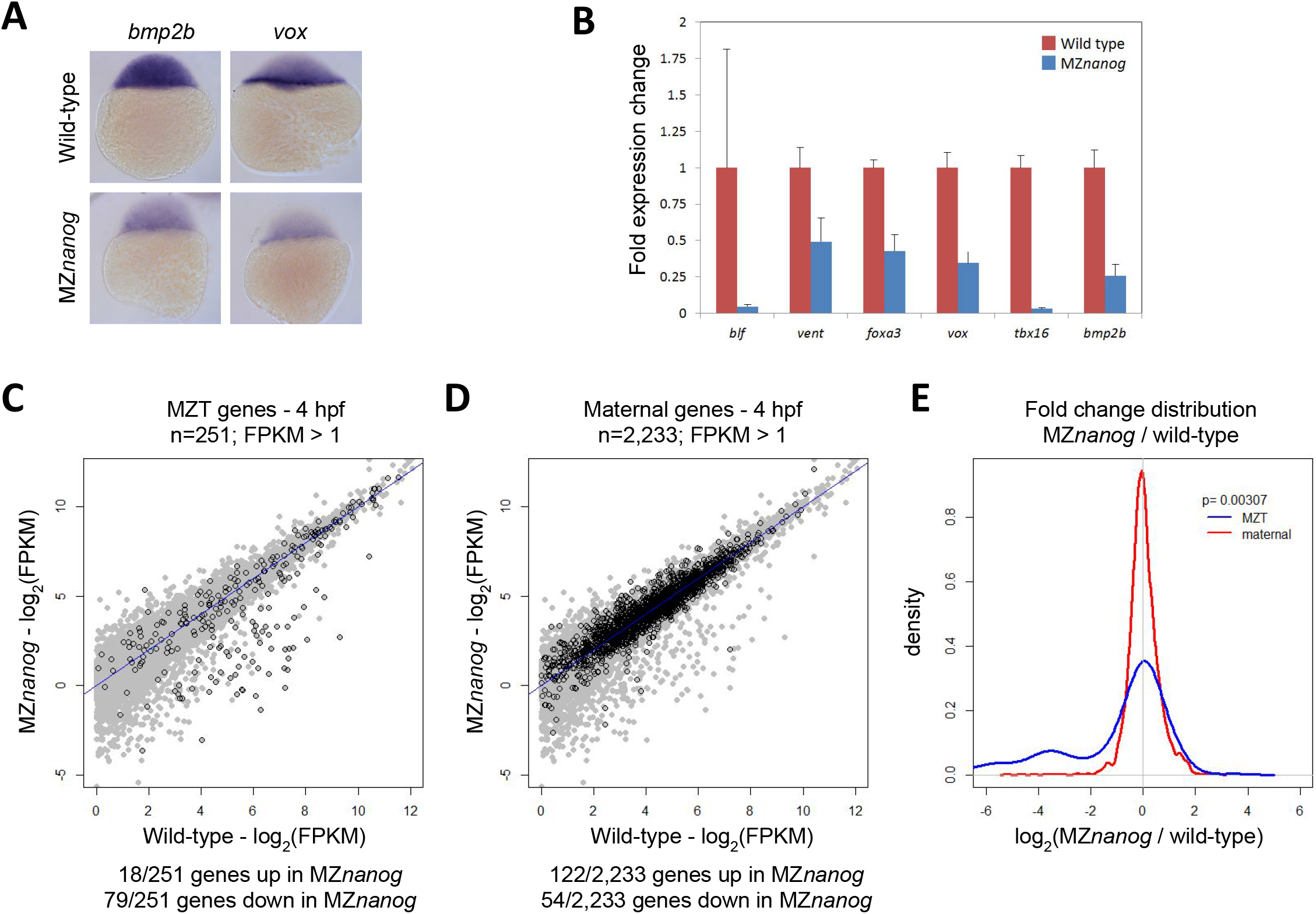
Defects in early zygotic gene expression in MZ*nanog* embryos. **A.** In situ hybridization for *bmp2b* and *vox* expression in wild-type and MZ*nanog* embryos at sphere stage. **B.** Fold expression change for the indicated genes comparing wild-type and MZ*nanog* at sphere stage using RT-qPCR. Error bars show standard deviation for three technical replicates. **C.** Differential expression of early zygotic genes, comparing wild-type and MZ*nanog* at sphere stage (4 hpf) using RNAseq. All genes are in grey, early zygotic genes in black were previously defined as the set of “first wave genes” (Lee et al. 2013). **D.** Differential expression of maternally provided genes, comparing wild-type and MZ*nanog* at sphere stage (4 hpf) using RNAseq. All genes are in grey, maternally provided genes in black were previously defined (Rabani et al. 2014). Only those genes with wild type mRNA expression > 1 FPKM are plotted. Genes are categorized as up- or down-regulated if their expression in MZ*nanog* differs more than two-fold from that in wild-type. **E.** Distribution of fold changes between wild-type and MZ*nanog* embryos for all early zygotic and maternal genes using a kernel density estimation. The displayed p-value comparing these two sets was calculated using a Student’s two-sided t-test.

### Expression of *nanog* mRNA in YSL precursors can rescue MZ*nanog* embryos

To clarify the embryonic and extraembryonic requirements for Nanog, we performed tissue-specific rescue experiments. We first repeated the previously described YSL rescue experiments using *mxtx2* mRNA injection (Xu et al. 2012). We deposited *mxtx2* mRNA in the precursor cells of the YSL by vegetal yolk injection at the 4-cell stage. Co-injection of *GFP* mRNA was used to verify that expression was limited to the YSL (**Figure 4A and E**). *Mxtx2* rescued the epiboly defect and yolk detachment in MZ*nanog* mutants (**Figure 4B**), and some rescued embryos developed differentiated tissues, including somites, notochord, eyes, brain, melanocytes, otoliths and blood. We then used the same approach to deposit *nanog* mRNA and *GFP* mRNA in the precursor cells of the YSL. Injection of *nanog* was sufficient to rescue the epiboly defects and yolk detachment in most MZ*nanog* mutants (**Figure 4C**). At 24 hours post-fertilization, a majority of mutants displayed axis rescue, tail elongation and differentiation of various cell types **(Figure 4D)**. Generally, rescue by *nanog* was more pronounced than by *mxtx2*. In contrast, injection of *nanog* mRNA into 1 of 16 cells did not result in any rescue of MZ*nanog* mutants (**Figure 4C and D**). These results indicate that the primary role of Nanog is to generate the YSL, and that YSL defects are a major cause of the embryonic phenotypes observed in MZ*nanog* mutants.

**Figure 4.**
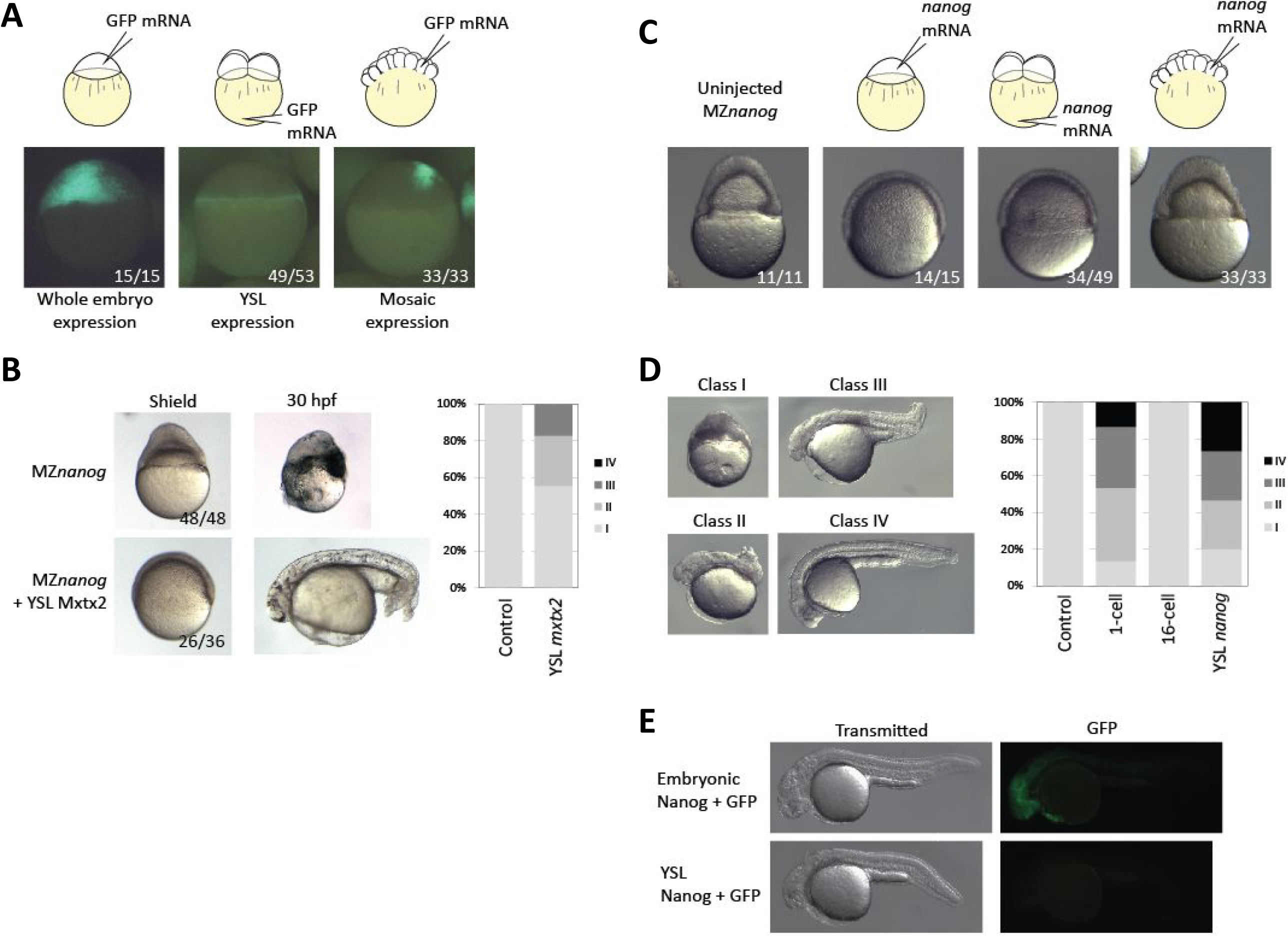
Rescue of MZ*nanog* embryos by YSL expression of *nanog* or *mxtx2* mRNA. **A.** MZ*nanog* embryos were injected with 33 pg GFP mRNA at 1-cell stage, 4-cell stage in vegetal yolk, or in 1 of the cells at 16-cell stage. At sphere stage, embryos were imaged and scored for spatial GFP expression. **B.** Control MZ*nanog* embryos, or embryos injected for YSL expression with 33 pg GFP mRNA and 50 pg *mxtx2* mRNA, were sorted for appropriate spatial GFP expression at sphere stage, scored as described in the Methods, and imaged at 75% epiboly stage and at 30 hpf. **C.** Control MZ*nanog* embryos, or embryos injected with 33 pg GFP mRNA and 25 pg *nanog* mRNA were sorted for appropriate spatial GFP expression at sphere stage, and scored and imaged at 75% epiboly stage. **D.** Embryos injected as indicated were scored at 24 hpf into categories as described in the Methods and shown in the panel. **E.** Representative rescued embryos (as indicated) imaged at 24 hpf for GFP expression in embryonic tissues.

### *Nanog* mutant cells can proliferate and differentiate in wild-type embryos

It has been proposed that Nanog is directly required in embryonic cells (Lee et al. 2013; Perez-Camps et al. 2016) but this assumption has not been tested. We therefore co-transplanted fluorescently labeled MZ*nanog* mutant and wild-type cells into wild-type hosts at sphere stage, and tracked their contributions to the host embryos. By 30 hpf, transplanted wild-type and MZ*nanog* mutant cells had proliferated to a similar extent in the host chimerae and contributed to many tissues, including brain, muscle and hatching gland (**Figure 5A**). To further test whether cells lacking Nanog can differentiate and contribute to tissues from all three germ layers, we used transgenic markers to follow donor cell differentiation, including *fli1*:GFP (vasculature, a derivative of the mesodermal germ layer), *acta*:GFP (trunk muscle, mesodermal germ layer), *islet*:GFP (trigeminal sensory neurons, ectodermal germ layer), and *sox17*:GFP (gastrointestinal tract, endodermal germ layer) (Sakaguchi et al. 2006; Higashijima et al. 2000; Higashijima et al. 1997; Lawson & Weinstein 2002). We transplanted cells from transgenic M*nanog* embryos into wild-type hosts and found that donor cells proliferated in the host embryo and activated each of the four marker transgenes (**Figure 5B**). These results reveal that zebrafish Nanog is not required for proliferation or differentiation of embryonic cells.

**Figure 5.**
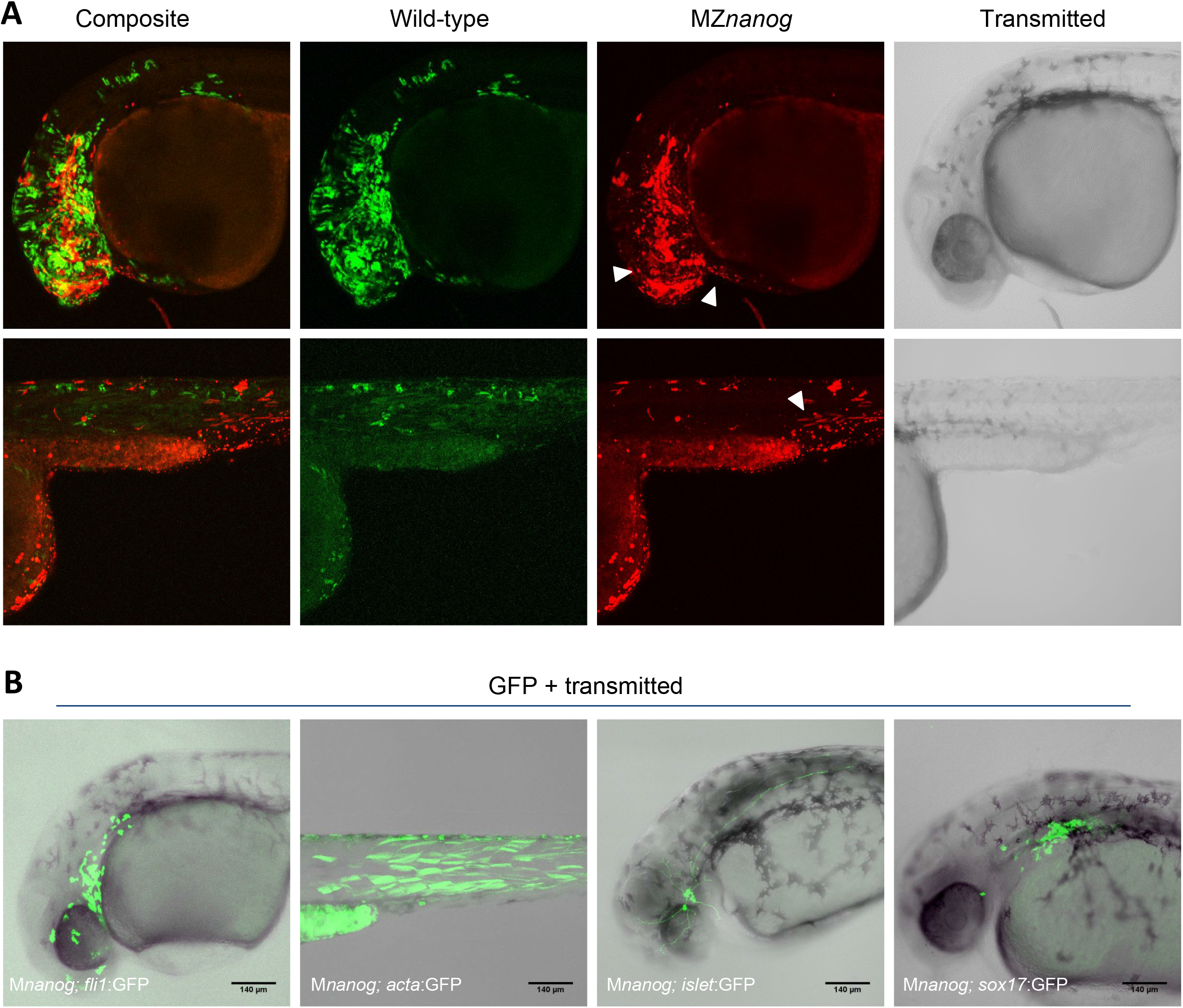
Transplantation of cells lacking Nanog into wild-type host embryos. **A.** Approximately 20 cells were transplanted from donor embryos injected with GFP mRNA (wild-type) or DsRed mRNA (MZ*nanog*) together into uninjected wild type host embryos. At 30 hpf, embryos were anaesthetized, mounted and imaged by confocal microscopy. Two representative embryos are pictured, with arrowheads indicating contributions to eye, hatching gland, and muscle fibers. **B.** Approximately 20 cells were transplanted from donor embryos (progeny of a Z*nanog* female crossed to a transgenic male) into uninjected wild type host embryos. At 30 hpf, embryos were anaesthetized, mounted and imaged by confocal microscopy. Representative embryos are displayed as maximum projections from a subset of a z-stack, with GFP transgene or tracer expression overlaid onto the transmitted light image for context.

## DISCUSSION

Our results support the conclusion that the primary role for zebrafish Nanog is in specification of the YSL (Xu et al. 2012) and that it has no essential autonomous functions in embryonic cells.

Three lines of evidence support an essential role of Nanog in YSL formation. First, both MZ*nanog* mutants (this study) and *nanog* morphants (Xu et al. 2012) lack expression of *mxtx2*, the master regulator of YSL maturation, as well as expression of several other YSL markers. Second, loss of Nanog blocks formation of the F-actin band within the YSL that powers epiboly (Xu et al. 2012) and leads to epiboly defects and embryo detachment (Xu et al. 2012; this study). Third, expression of *nanog* mRNA (this study) or *mxtx2* mRNA (Xu et al. 2012; this study) in YSL precursors partially rescues epiboly and embryonic patterning caused by the loss of Nanog. These observations strengthen and extend the previously reached conclusion (Xu et al. 2012) that Nanog is essential for YSL formation.

Two lines of evidence indicate that Nanog is not directly required in embryonic cells. First, *nanog* (this study) or *mxtx2* (Xu et al. 2012; this study) expression in YSL precursors is sufficient to rescue many aspects of the phenotype, including epiboly, embryo attachment and embryonic cell differentiation. Second, embryonic cells that lack Nanog and are transplanted into wild-type hosts proliferate and differentiate into derivatives of all germ layers (this study) and germ cells (unpublished results). These observations indicate that Nanog has no essential autonomous role in embryonic cell differentiation.

Although our results cast doubt on the previously held assumption that Nanog is directly required in embryonic cells for zygotic gene activation (Lee et al. 2013), we did find that MZ*nanog* embryos exhibit embryonic gene expression defects during MZT. 79 genes expressed at sphere stage displayed a two-fold or greater reduction of transcript levels in MZ*nanog* mutants. Many of these genes encode components of developmental signaling pathways. The gene set reduced in *nanog* mutants broadly overlapped with genes downregulated in *nanog* morphants (Lee et al. 2013). Thus, previously observed gene regulation defects (Xu et al. 2012; Lee et al. 2013) were not simply a consequence of delayed development in morphants. Moreover, maternally provided *nanog* is present throughout the embryo (Figure 1 and Xu et al. 2012), and Nanog binds promoters of many developmental regulators in the early embryo (Xu et al. 2012; Lee et al. 2013; Leichsenring et al. 2013; Bogdanovic et al. 2012). Although Nanog is required in in YSL formation but not embryonic cell differentiation, these results might suggest a potential contribution of Nanog to the expression of some genes during MZT.

How then can the seemingly contradictory observations on the roles of Nanog be reconciled? We suggest the following model. Nanog is primarily required for the formation of the YSL and the activation of the YSL gene expression program. These genes include patterning signals that in turn regulate gene expression in embryonic cells. Nanog is not required for zygotic genome activation but binds in conjunction with other pluripotency factors to cis-regulatory regions of embryonic genes and makes non-essential contributions to their transcription. In this scenario zebrafish Nanog primarily acts extraembryonically and its embryonic requirements are similar to those in mammals - involved in but not essential for the acquisition and maintenance of pluripotency.

## MATERIALS AND METHODS

### Animal care

All vertebrate animal work was performed at the facilities of Harvard University, Faculty of Arts & Sciences (HU/FAS). The HU/FAS animal care and use program maintains full AAALAC accreditation, is assured with OLAW (A3593-01), and is currently registered with the USDA. This study was approved by the Harvard University/Faculty of Arts & Sciences Standing Committee on the Use of Animals in Research & Teaching under Protocol 25–08.

### Generation of the *nanog* mutants

A TALEN pair targeting the first exon of *nanog* (TALEN L: TCCCGAATCTGAGCTGGC, TALEN R: TGTGACCCCGCCGGAGTGT) was generated using the TALE Toolbox (Sanjana et al. 2012). Wild type embryos were injected at the 1-cell stage with 450 pg TALEN pair mRNA. Mutations were verified in injected embryos, and from clutches of outcrossed putative founder adults, by PCR from genomic DNA, followed by T7 Endonuclease I assay (NEB). Mutations within individual founder fish were identified by cloning PCR products followed by Sanger sequencing. An allele containing a 7 bp deletion was isolated and used for all further experiments. These fish were genotyped using a PCR strategy - two allele-specific primers, in combination with a constant primer, separately identify wild type and mutant alleles. Allele description, primer and vector sequences can be found in **Table S2**.

### Molecular cloning and in situ hybridization

Total RNA was isolated from embryos using EZNA Total RNA kits (Omega Biotek). cDNA was generated using iScript cDNA Synthesis kit (Bio-Rad). Open reading frames for *nanog* and *mxtx2* were amplified by PCR from embryonic cDNA and cloned into the pCS2 vector for generation of mRNA and probes. Open reading frames for *vox* and *bmp2b* were amplified by PCR and cloned using the Strataclone kit (Agilent) for generation of probes. mRNAs were transcribed using mMessage mMachine kits (Ambion). Antisense probes for in situ hybridization were transcribed using DIG RNA labeling kit (Roche). All RNAs were purified using EZNA Total RNA kits (Omega Biotek). In situ hybridization was completed as previously described (Thisse & Thisse 2008); stained embryos were cleared using BB:BA and imaged with a Zeiss Axio Imager.Z1 microscope. Vector sequences can be found in **Table S2**.

### RT-qPCR

Total RNA and cDNA were generated as above. qPCR was conducted using iTaq (Bio-Rad) on a CFX96 (Bio-Rad). Primer sequences can be found in **Table S2**.

### RNAseq

Total RNA was isolated from MZ*nanog* and wild type embryos at sphere and shield stages (n=40 embryos each condition) following a previously published protocol (Pauli et al. 2012). RNA quality was confirmed by Bioanalyzer (Agilent). RNAseq libraries were generated using the TruSeq RNA Library Prep Kit v2 (Illumina) and sequenced on a HiSeq 2500, generating single-end 51 bp reads. Reads were aligned for each sample using TopHat v2.0.13 (Trapnell et al. 2009) with the following command “tophat -o <output directory> -p 16 –no-novel-juncs -G <gene table> <Bowtie2 genome index> <fastq reads>”. Transcript abundance and differential expression were determined using Cufflinks v2.2.1 (Trapnell et al. 2012) with the following command for each developmental stage “cuffdiff -p 16 -b <genome.fa -u -L <labels> -o <output directory> <gene table> <wild type aligned reads .bam file> <mutant aligned reads .bam file>”. Differential gene expression plots were generated in Rstudio.

### YSL expression and phenotypic scoring

YSL expression was performed through injection of mRNAs at the 4-cell stage into the vegetal yolk (Xu et al. 2012). Embryos with expression restricted to the YSL were scored and sorted at sphere stage using fluorescence from co-injected GFP mRNA. Phenotypes for all injected embryos were scored during gastrulation and at 24-30 hpf. At 24-30 hpf, classification used the following category definitions: Class I - no rescue, ball of necrotic cells or exploded; Class II - axis rescue; Class III, axis rescue and tail extension; Class IV, similar to wild-type.

### Transplantation and imaging

Non-transplanted embryos were anaesthetized when necessary, mounted in 3% methylcellulose, and imaged with a Leica MZ 16 F microscope. For transplantation, donor embryos were injected, when appropriate, with either GFP mRNA or DsRed mRNA (50 pg each). Approximately 20-40 cells from a donor embryo were transplanted at sphere stage to a host embryo. Transplant host embryos were screened for fluorescence with a Leica MZ 16 F microscope, then mounted in 1% low melt agarose and imaged with a Zeiss Pascal confocal microscope.

## ACKNOWLEDGEMENTS

Thanks to members of the Zon lab for the *fli1*:GFP line and helpful discussions, Michal Rabani, Jeff Farrell, Katherine Rogers, Megan Norris and Andrea Pauli for technical assistance, and the Harvard Bauer Core Facility for sequencing.

## AUTHOR CONTRIBUTIONS

J.A.G. and A.F.S. conceived of the project and designed experiments. J.A.G., with assistance from K.O., performed and completed all experiments. J.A.G. and A.F.S. interpreted results and wrote the manuscript.

## DATA AVAILABILITY

RNAseq data has been uploaded to GEO under accession number GSE89245.

